# CIGB-300 synthetic peptide, an antagonist of CK2 kinase activity, as a treatment for Covid-19. A computational biology approach

**DOI:** 10.1101/2021.07.26.453805

**Authors:** Jamilet Miranda, Ricardo Bringas, Jorge Fernández-de-Cossio, Yasser Perera

## Abstract

Drug repositioning became the first choice for treating Covid-19 patients due to the urgent need to deal with the pandemic. Similarities in the hijacking mechanisms used by SARS-CoV-2 and several type of cancer, suggest the repurposing of cancer drugs to treat Covid-19. CK2 kinase antagonists have been proposed for the treatment of cancer. A recent study in cells infected with SARS-CoV-2 virus found a significant CK2 kinase activity, and the use of a CK2 inhibitor showed antiviral responses. CIGB-300, originally designed as an anticancer peptide, is an antagonist of CK2 kinase activity that binds to CK2 phospho-acceptor sites. Recent preliminary results show an antiviral activity of CIGB-300 versus a surrogate model of coronavirus. Here we present a computational biology study that provides evidences at the molecular level of how CIGB-300 might interfere with SARS-CoV-2 life cycle inside infected human cells. First, from SARS-CoV studies, we infer the potential incidence of CIGB-300 in SARS-CoV-2 interference on immune response. Next, from the analysis of multiple Omics data, we propose the action of CIGB-300 since early stage of viral infections perturbing the virus hijacking of RNA splicing machinery. It was also predicted the interference of CIGB-300 in virus-host interactions responsible for the high infectivity and the particular immune response to SARS-CoV-2 infection. Further, we provide evidences of CIGB-300 attenuation of phenotypes related to muscle, bleeding, coagulation and respiratory disorders.

## Introduction

SARS-CoV-2 currently spread world-wide showing high infectivity and transmissibility. Due to the urgency of finding effective therapeutics treatments in the shortest possible time, drug repurposing emerged as the first option [1, 2]. The huge amount of data so far generated permit the re-consideration of drugs already evaluated for other diseases, which might have advanced toxicological, preclinical and/or clinical studies.

Since the genomic sequence of SARS-CoV-2 was made available in January 2020 [3], diversity of technique and laboratory models has been profusely applied to the study of SARS-CoV-2 replication and infectivity.

Blanco-Mello et al. [4] performed RNA-seq experiments from polyA RNAs isolated from infected cells. They found a diminished transcriptional response of type I/III interferon-induced genes and, concurrently, a significant increase of chemokines and IL6, which ground their suggestion to evaluate FDA-approved drugs with immunomodulating properties that could be rapidly implemented in clinical protocols.

Various mass-spectrometry studies made important contribution to the comprehension of SARS-CoV-2 life cycle. Gordon et al. [5] using affinity-purification mass spectrometry (AP-MS) identified the sets of human proteins that physically interact with each of 26 viral proteins that they previously individually cloned in human cells derived from kidney HEK-293T/17. A total of 332 physical interactions between SARS-CoV-2 and human proteins were identified. When the expressions of all interacting human proteins in 29 different tissues were analyzed, lung was identified as the one with the highest expression levels. GO enrichment analysis was performed to the set of human interactors of each viral protein cloned. For each of the 26 sets, the major overrepresented biological processes included Nuclear Transport, Ribonucleoprotein Complex Biogenesis and Cellular Component Disassembly. Host proteins involved in innate immune response were targeted by viral proteins nsp13, nsp15 and orf9b while proteins from the Nf-kB pathway were targeted by nsp13 and orf9c. The most relevant host proteins targeted by known drugs were identified from this analysis. Next, they demonstrated the capacity of some of these drugs to reduce viral infectivity. The study of Gordon et al. [5] is of outstanding relevance for the understanding of the mechanism used by SARS-CoV-2 to improve its infectivity and to avoid a strong immune response. It also provides valuable information for repurposing of existing drugs.

The role of kinases in the course of viral infection was addressed by Bouhaddou et al. [6], who carried a quantitative mass spectrometry-based phosphoproteomics study in Vero E6 cells infected by SARS-CoV-2. Casein kinase II (CK2) and p38 MAP kinases were significantly activated while mitotic kinases were shutdown. A relevant role of CK2 in induced filopodia protrusions during viral infection was evidenced in association with viral capsid protein N. Both CK2 and protein N were co-localized in filopodia protrusions which were significantly longer and more branched than in control cells. The authors suggest that N protein may control CK2 activity and regulate cytoskeleton elements in filopodia. Although the role of CK2 in viral infections is not new, it is remarkable the level of upregulation of CK2 activity as consequence of SARS-CoV-2 infection [6]. The strong antiviral activity of Silmitasertib (CX-4945), a CK2 inhibitor, suggests this kinase as an attractive target for treating Covid-19 patients. Ongoing clinical trial of CX-4945 is evaluating its clinical benefits and anti-viral activities in moderate COVID-19 patients (https://clinicaltrials.gov/ct2/show/study/NCT04663737).

CIGB-300 is a synthetic peptide designed to bind the phospho-acceptor motif of CK2 substrates, interfering the phosphorylation of serine/threonine residues by CK2. Clinical use of CIGB-300 have confirmed it safety and tolerability when administered intravenously showing clinical efficacy in cancer patients [7, 8].

In a phosphoproteomic experiment in NCI-H125 cells, Perera et al. [9] identified CK2 phospho-acceptors peptides that are significantly inhibited by CIGB-300. They found for the first time, that CIGB-300 binds CK2α subunit and impairs CK2α2β2 holoenzyme enzymatic activity. By contrast, phosphorylation of CK2β subunit, which contains itself a consensus CK2 phosphorylation motif, was not influenced by CIGB-300. Additionally, Perera et al. [10] identified nucleophosmin (B23) as a major target of CIGB-300. Of note, Nouri et al. [11] further reported the binding of CIGB-300 to B23 oligomerization domain. This interaction blocks the association of B23 to Rev and US11 proteins, two functionally homologous proteins from HIV and HSV viruses respectively. Cells treated with CIGB-300 showed a significant reduction of virus production suggesting B23 as an attractive target for antiviral drugs. Lobaina and Perera [12] also proposed B23 as a potential target in antiviral therapies.

With this background, CIGB-300 was tested for its safety and clinical benefits in Covid-19 patients in a phase I/II clinical trial [13]. It reduced the number of pulmonary lesions among treated individuals. Additionally, CIGB-300 antiviral effect on MDBK cell infected with bovine coronavirus (BCoV) Mebus was explored [14]. CIGB-300 inhibited the cytopathic effect and reduced the viral protein accumulation in the cytoplasm. Physical interaction of CIGB-300 with BCoV nucleocapsid protein(N) was also revealed. Functional enrichment found cytoskeleton reorganization and protein folding as the major disturbed biological processes.

Here we present an *in silico* analysis of SARS-CoV and SARS-CoV-2 viral infection. We performed a multi-omics integrative analysis of SARS-CoV-2 infection of human cell lines that combines functional enrichment and network representation. At the level of phosphorylation sites, we integrated data from four different phosphoproteomics studies on SARS-Cov-2 infection [6,15–17] and one study on CIGB-300 inhibition of kinase activity [9]. We identified biological processes and virus activated phosphosites at different times after viral infection that can be interfered by CIGB-300. Our results are consistent with the benefits already evidenced of CIGB-300 treatment in Covid-19 patients.

## Results and Discussion

### Inferring the potential effect of CIGB-300 treatment on SARS-CoV-2 virus infection based in previous results

#### CIGB-300 could alter N-protein localization and its RNA binding capacity

Coronavirus nucleocapsid N proteins play an essential role in virus cell cycle; its dimerization and binding to the viral genomic RNA is the first step for virion particle assembly. N protein plays an important role in viral genomic RNA synthesis [18] and have been also implicated in the inhibition of type I interferon signaling pathway [19].

A sequence alignment of N protein from SARS-CoV and SARS-CoV-2 viruses show a high similarity (Fig 1). N Protein consists of two structural domains (Fig 1): the N-terminal RNA-binding domain (RBD) (residues 41–186), and the C-terminal dimerization domain (residues 258-361). The rest of the protein is highly disordered [20].

**Figure 1:**
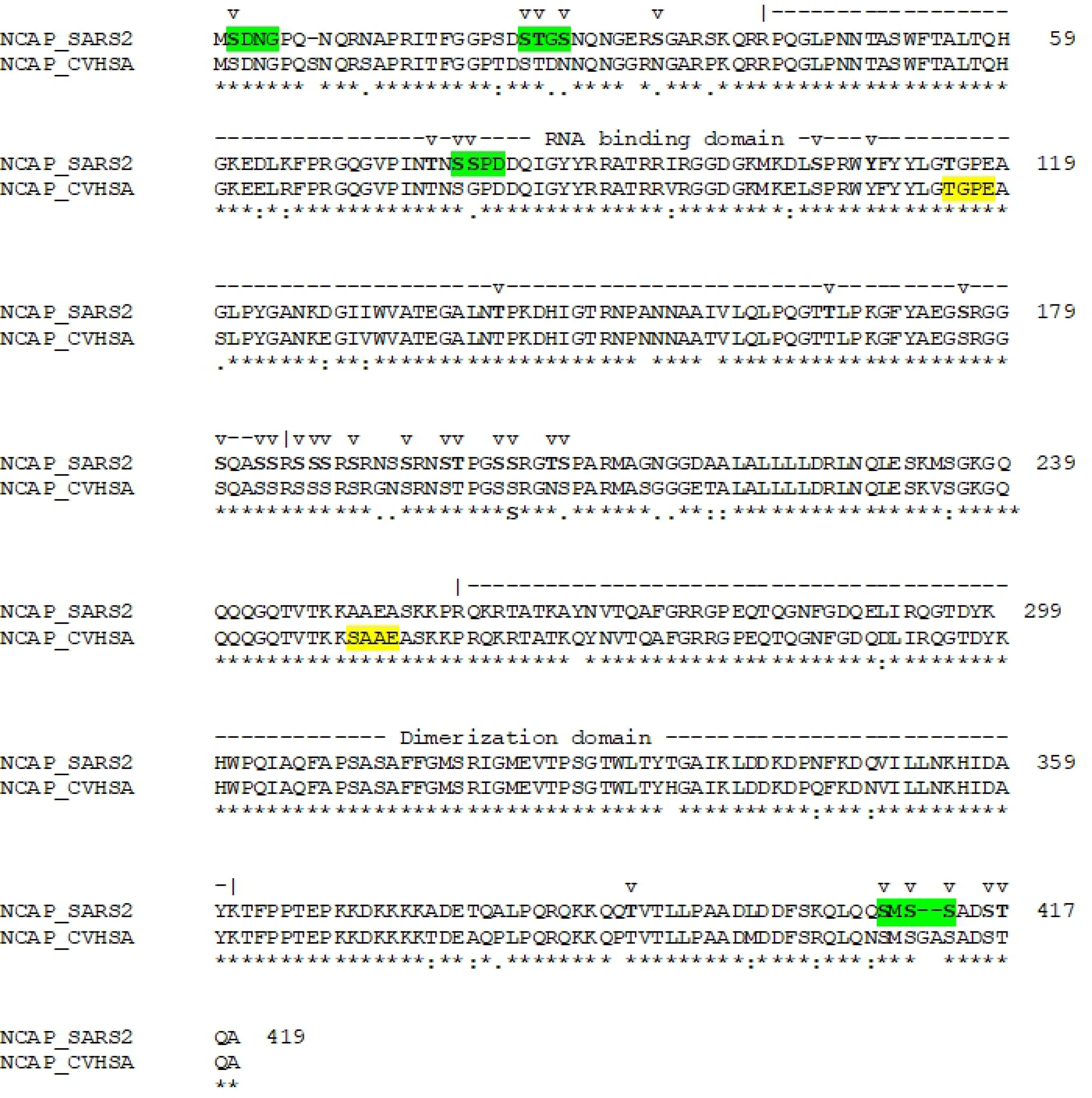
Sequence alignment of nucleocapside proteins of SARS-CoV-2 (NCAP_SARS2) and SARS-CoV (NCAP_CVHSA) viruses. Numbers at the right end indicate amino acids position in the SARS-CoV-2 protein. Above the aligned sequences the RNA-binding (residues 41-186, according to UniProt annotation) and Dimerization (residues 258-361) domains are delimited. v-above aminoacid residues indicates phosphorylation sites identified in the works of Davidson et al. [21], Bouhaddou et al. [6], Klann et al. [16] and Hekman et al. [15]. Highlighted are the segment between a CK2 phospho-acceptor site and position +3, in yellow those *in silico* predicted by Surjit et al. [22] and in green those experimentally validated by Davidson et al. [21].

Surjit et al. [22] predicted SARS-CoV N-protein to be heavily phosphorylated. Thr116 and Ser251 were noted as putative phospho-acceptors for CK2 (see Fig 1), though neither has been corroborated experimentally. We collected a total of 33 phosphorylation sites for SARS-CoV-2 N protein (see Fig 1 and S1 Table) from four recent mass spectrometry studies [6,15,16,21] reported two putative CK2 sites at Ser2 and Ser78. Hekman et al. [15] found Ser23 and Ser410 to be phosphorylated by CK2.

Bouhaddou et al. [6] analyzed the impact of phosphorylation in the N-protein surface charges by a 3D structural model of the RNA-binding domain. These changes may modulate the function of N-protein by regulating its RNA binding capacity. One of the phosphorylation sites responsible for these charge changes is the Ser78 (see Fig 1), a CK2 phospho-acceptor site according to Davison et al. [21]. Binding of CIGB-300 to Ser78 would interfere with N-protein RNA binding ability. On the other hand, N protein is reported to be mainly located in the cytoplasm [22–24]. However, a localization analysis of N-expressing cells treated with four different phosphorylation inhibitors found a significant fraction of N protein localized in the nucleus of cells treated with CDK or CK2 inhibitors [22]. Additionally, in cell infected by BCoV, CIGB-300 bound N protein, downregulated its expression and significantly reduced the accumulation of viral proteins in the cytoplasm [14].

Bouhaddou et al. [6] found CDK activity to be significantly reduced by SARS-CoV-2 infection while CK2 activity is significantly increased. Consequently, inhibition by CIGB-300 of N protein phosphorylation sites may alter, at least in part, its cytoplasmic localization. Hence, the use of CIGB-300 in Covid-19 patients would interfere the N protein role in viral cell cycle in infected cells as its function in particle assembly happens in cytoplasm.

#### CIGB-300 could bind ORF6 C-terminus and restore IFNs signaling

One important element of innate immune response to virus infections is the activation of antiviral genes as a consequence of interferon production. After activation of receptors by type I interferons, STAT1 is phosphorylated and forms a complex with STAT2 and IRF9 [25]. This complex exposes a nuclear localization signal (NLS) that is bound by KPNA1, and as a last step before entering the nucleus KPNB1 binds KPNA1 and chaperons the complex through the nuclear pore (Fig 2A) [26].

**Figure 2:**
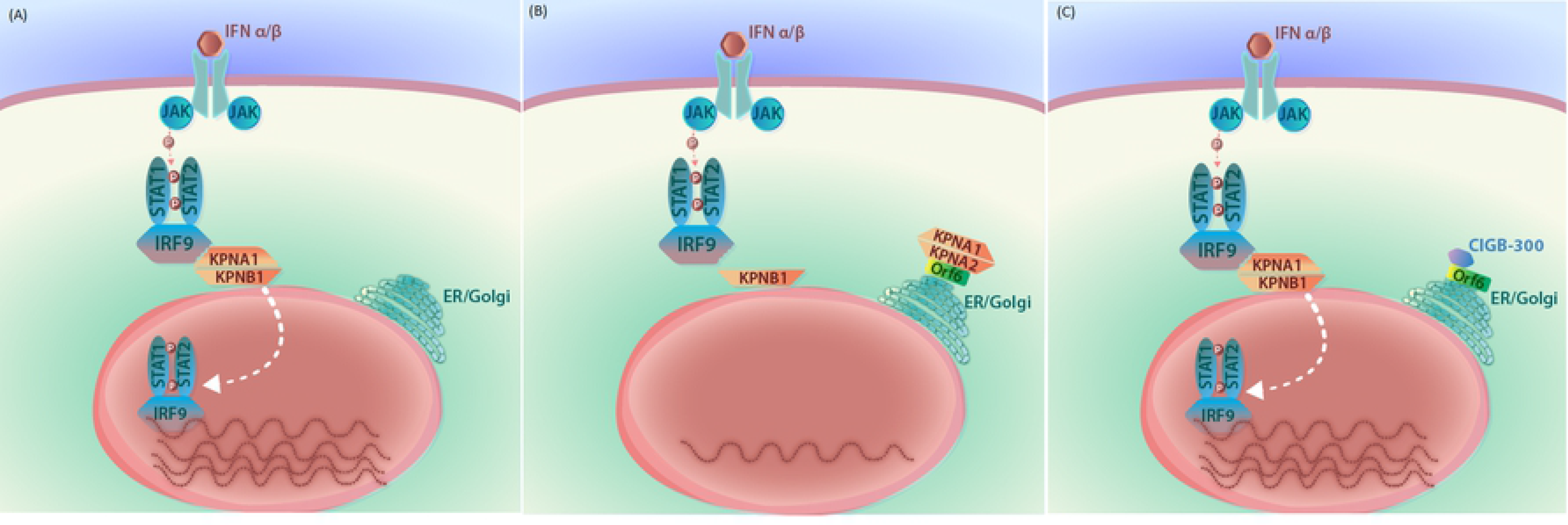
Interference of IFN signaling by Orf6 and the role of CIGB-300. A. Interferon binding to receptor induces STAT1 phosphorylation and formation of a complex with STAT2 and IRF9. KPNA1 binds the complex, KPNB1 binds KPNA1 and chaperons the complex through the nuclear pore. B. Orf6 retains KPNA1 and KPNA2 in the ER/Golgi membrane and the transport of STAT complex to the nucleus interrupted. C. CIGB-300 blocks the interaction of Orf6 with KPNA2 and the transport of STAT complex to the nucleus is restored.

Several groups have attributed an immune response antagonistic effect to Orf6 protein [19,26,27]. In SARS-CoV experiments, Frieman et al. [26] reported that Orf6 interferes with host immune response by antagonizing STAT1 function. Orf6 binds karyopherin alpha 2 (KPNA2) and retains it in the ER/Golgi membrane. KPNB1 is also retained as it binds KPNA2. In this way, the chaperon function of KPNB1 through the nuclear pore is interfered, and STAT1 signaling is interrupted (Fig 2B).

Frieman et al. [26] also found that the C-terminal 10 amino acids of SARS-CoV Orf6 are responsible for KPNA2 binding. In Fig 3 we show the residues of Orf6 involved in the SARS-CoV mutants they generated, Orf6_49-53Ala_, Orf6_54-58Ala_ and Orf6_59-63Ala_ (author’s nomenclature), by replacing amino acids 49-53, 54-58 and 59-63 with alanines, respectively. The last two mutants, Orf6_54-58Ala_ and Orf6_59-63Ala_, comprising the ten C-terminal amino acids, did not retain KPNA2 and as consequence, STAT1 function was unaffected. The first mutant Orf6_49-53Ala_ was still able to retain KPNA2. So, the last ten aminoacids were responsible for KPNA2 binding and, as consequence, for KPNB1 recruitment.

**Figure 3:**
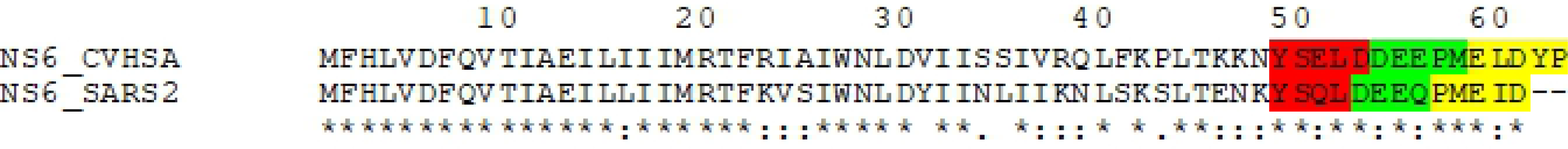
Sequence alignment of Orf6 proteins of SARS-CoV-2 (NS6_CVHSA) and SARS-CoV-2 (NS6_SARS2) viruses. Highlighted in red, green and yellow are the residues that were replaced by alanines in the mutants generated by Frieman et al. [26] and Lei et al. [28].

Recently Lei et al. [28] carried a similar mutation study of SARS-CoV-2 Orf6 protein. They generated three different mutants; M1, M2 and M3 (author’s nomenclature); by replacing aminoacids 49-52 (YSQL), 53-56 (DEEQ) and 57-61 (PMEID) by alanines, respectively (Orf6 of SARS-CoV-2 lacks the last two amino acids present in SARS-CoV protein). As expected, they obtained similar results: mutant M1 perturbs interferon stimulation as the wild type does, while mutants M2 and M3 lack the inhibitory effect.

In Fig 3 we show an alignment of Orf6 protein sequences from SARS-CoV and SARS-CoV-2 viruses. The region between amino acids 50-53 with the sequence SELD in SARS-CoV protein and sequence SQLD in SARS-CoV-2, both match the CK2 substrate motif. Additionally, this site in SARS-CoV-2 was experimentally found to be phosphorylated, and predicted by computer analysis to be a phospho-acceptor site of CK2 [16]. Ser50, as a CK2 phospho-acceptor site, could be bound by CIGB-300.

Mutant M2 of Lei et al. [28] include Asp53 residue at position +3 relative to Ser50, and this position is known to be important for the recognition of CK2. Therefore, we strongly suggest that the possible binding of CIGB-300 to this phospho-acceptor motif would interfere the interaction of Orf6 C-terminus with KPNA2; avoiding its retention in the ER/Golgi membrane, without interfering KPNA2 chaperon activity of carrying STAT1 complex to the nucleus (Fig 2C). In this regard, CIGB-300 could exhibit an effect that other CK2 antagonists that target CK2 won’t.

#### Interfering NUP98 hijacking by CIGB-300 via interaction with Orf6 C-terminus

We analyzed proteomic expression data from Bojkova et al. [29] and found B23 exhibit the highest positive correlation with the expression profile of viral proteins (S1 Fig). SARS-CoV virus N protein was found to interact with B23 protein [30]. Despite that, Gordon et al. [5] did not reported a direct interaction of B23 with viral proteins. Looking for indirect interactions, we intersected the interactors of B23 with the 322 proteins found by Gordon et al. [5] to interact with viral proteins. A total of 21 host proteins resulted from this intersection, among which Nuclear Pore Complex protein 98 (NUP98) shown up as the only one that interact with Orf6, the viral protein with the highest expression correlation to B23.

Bouhaddou et al. [6] determined that phosphorylation at Ser888 of NUP98 increased during viral infection. The sequence around Ser888 is D**S**DEEE, which fulfills the phospho-acceptor motif of CK2. Additionally, Franchin et al. [31] found the phosphorylation of Ser888 to be altered by a CK2 inhibitor (according to data downloaded from PhosphositePlus web site). NUP98 is part of the Nuclear Pore Complex, responsible for the transport of biomolecules between the nucleus and cytoplasm. Bouhaddou et al. [6] suggested that the SARS-CoV-2 infection-induced phosphorylation of NUP98 may prevent export of mRNAs through the nuclear pore, a similar mechanism to those used by other viruses to increase the translation of viral RNA in the cytoplasm. Binding of CIGB-300 to Ser888 phospho-acceptor site of NUP98 could prevent its phosphorylation and restore host mRNA translocation to cytoplasm.

Also, Gordon et al. [5] found that Met58 and acidic residues Glu55, Glu59 and Asp61 are highly conserved in Orf6 homologs and are part of a putative NUP98/RAE binding motif. Miorin et al. [32] found that SARS-CoV-2 infection blocks the nuclear translocation of STAT1 and STAT2. Orf6 exerts this anti IFN-I activity by hijacking NUP98. Orf6 directly interact with NUP98 at the Nuclear Pore Complex(NPC) via its C-terminal end. A Met58Arg mutant in Orf6 C-terminal region impairs this interaction and abolish the IFN-I antagonistic effect [32].

The Orf6 interactions with KPNA2 and NUP98 have been both reported to interfere with IFN signaling. In both cases the C-terminal domain of Orf6 was responsible for the interaction, mutations in this region abolished the anti-IFN activity. The binding of CIGB-300 to the CK2 phospho-acceptor site Ser50 in Orf6 could impair the interaction with both KPNA2 and NUP98 and in some extend restore IFN signaling.

### CIGB-300 downregulate host proteins phosphosites consistently activated by SARS-CoV-2

We now compare the phosphoproteomics studies of SARS-CoV-2 infection in Vero E6 [6], Caco-2 [16], iAT2 [15] and A549 [17] cell lines with that of Perera et al. [9] on CIGB-300 kinase antagonistic effect in H125.

First, we combined results of the four studies at the level of phosphorylation sites and found a total of 8642 different sites that were upregulated in at least one of the studies (Venn diagram in Fig 4A). As noted by Hekman et al. [15], there are few proteins differentially regulated that coincide in all the four studies. Indeed, we found only six phosphosites that were upregulated by SARS-CoV-2 infection in the four cell lines.

**Figure 4:**
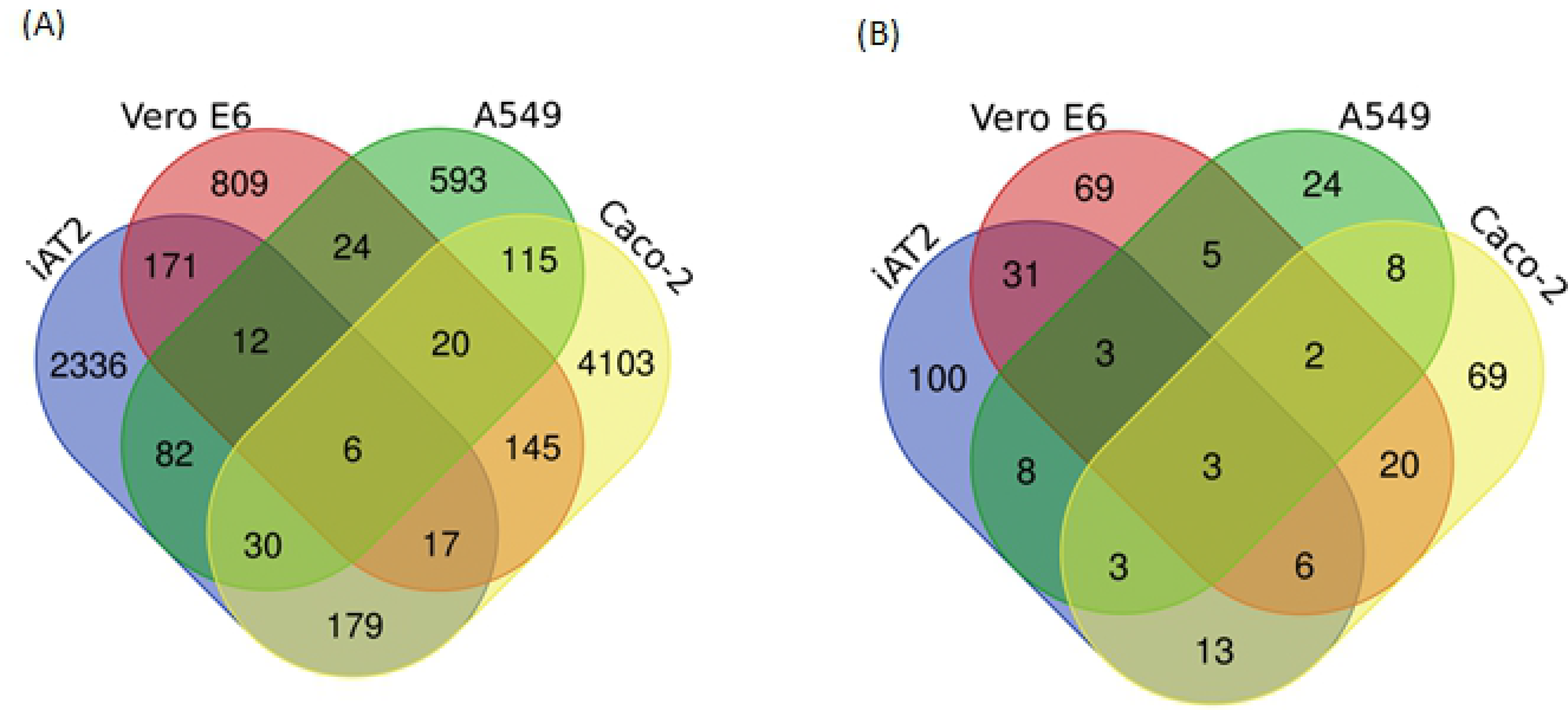
Results of four phosphoproteomics studies on SARS-CoV-infection in iAt2, Vero E6, A549 and Caco-2 cell lines and one study on CIGB-300 treatment effect on phosphorylation in H125 cell line. A. Venn diagram showing unique phosphorylation sites identified as UP regulated in A549, VeroE6, iAT2 and Caco-2 cells. B. Venn diagram showing unique phosphorylation sites identified as UP regulated in A549, VeroE6, iAT2 and Caco-2 cells and DOWN regulated by the action of CIGB-300.

Next, we intersected the data on SARS-CoV-2 infection with that of phosphorylation sites down regulated by the treatment of H125 cell line with CIGB-300, resulting in a total of 364 sites (see Fig 4B). Of the six sites that were found upregulated in the four phosphoproteomics studies, half were downregulated by CIGB-300. These three sites, MATR3_S188, SQSTM1_S272 and DIDO1_S1456, have in common to have tens of Phosphorylation sites. We envisage that these three phospho-acceptor sites, can be targeted by a CIGB-300 treatment in Covid-19 patients.

MATR3 is a nuclear matrix protein with 36 phosphosites according to UniProt annotations. MATR3 plays multiple functions in DNA/RNA processing, it contains two RNA recognition Motifs and two Zinc Finger domains. It was proposed to stabilize mRNA species, to play a role in the regulation of DNA virus-mediated innate immune response [33] and to be associated to splicing regulation [34]. In HIV-infected cells, Sarracino et al. [35] found MATR3 to be essential for RNA processing. MATR3 phoshorylation was found to greatly enhance its DNA binding ability [36, 37]. It is well documented its implications in Amyotrophic lateral sclerosis [38], a disease causing muscle weakness and respiratory failures, symptoms common in Covid-19 patients. CIGB-300 interference on virus-infection induced phosphorylation of MATR3 may play a role diminishing its effects in immune response attenuation and its implications in viral RNA processing.

SQSTM1 exhibit several phosphorylation sites, of these, Ser272 is the only one significantly activated by Sars-CoV-2 in the four phosphoproteomic studies. Zhang et al. [39] found that phosphorylation of SQSTM1 at Thr269 and Ser272 by MAPK13 promotes the microaggregates transport to the microtubule organizing center (MTOC) to form aggresomes which are later degraded through autophagy. Gao et al. [40] also showed that SQSTM1 phosphorylation increases its ability to sequester ubiquitinated proteins into aggresomes playing an important role in aggresome formation. Stukalov et al. [17] revealed significant reduction of autophagy flux by ORF3 which combined with the augmented microaggregates transport due to SQSTM1 phosphorylation conduces to the accumulation of aggresomes.

Several studies have reported the role of SQSTM1 accumulation and aggresome formation in lung related diseases. Tran et al. [41] demonstrated the role of aggresome formation induced by cigarette smoke in chronic obstructive pulmonary disease (COPD). They found a significant higher accumulation of SQSTM1 in smokers as compared to nonsmokers, and an increased severity of COPD. Wu et al. [42] found that the accumulation of SQSTM1 plays a critical role in airway inflammation induced by nanoparticles.

Cystic fibrosis (CF), is caused by mutations in the gene encoding the cystic fibrosis transmembrane conductance regulator (CFTR), which results in defective autophagy, causing the accumulation of CFTR containing aggregates [43]. SQSTM1 knockdown favoured the clearance of defective CFTR aggregates [44].

Inhibition by CIGB-300 of SQSTM1 phosphorylation at Ser272 may reduce the accumulation of aggresomes and this way attenuates lung inflammation and fibrosis induced by viral infection.

DIDO1 (death inducer-obliterator 1 or death-associated transcription factor DATF1) is a protein involved in apoptosis and have been also implicated in the progression of several type of cancer [45–48]. DIDO1 possess 92 phosphosites, according to data we downloaded from Phosphosite. Of these sites we found 16 that match the CK2 phospho-acceptor motif described by Pinna [49]. It is not clear the implications of DIDO1 in the course of viral infection, but it is known the induction of apoptosis by viral proteins and then DIDO1 may be activated by apoptosis pathway through phosphorylation. CIGB-300 may interfere this activation.

#### CIGB-300 at early Stage of SARS_CoV-2 Infection

We examine kinase activity from the earliest stages of the viral infection by analyzing phosphoproteomics data of Bouhaddou et al. [6] at 2h and 4h time points, and from Hekman et al. [15], at 1h and 3h time points.

GSEA analysis with proteins sets ranked by phosphorylation changes was performed to identify enriched REACTOME pathways.

After one hour of infection we observed a clear initial inhibition of host protein synthesis machinery, reflected in the inactivation of several phosphorylation sites of proteins involved in RNA metabolism events, such as “mRNA Splicing” and the “Pre-processing of capped intron containing mRNA” (Fig 5). This inactivation is immediately reverted by the activation of these same biological events at 2h and 3h. Fig 5 show the list of most significant pathways at each time point. The phosphosites listed are of those belonging to proteins from the core enrichment set of each enriched pathway and that were also identified by Perera et al. [9] to be inactivated by CIGB-300. Of those, SRSF1_S199 was the only site to be up-regulated at 2h and 3h. SRSF1_S201 was up-regulated at 3h as well as some other SRSF’s proteins phosphosites (Fig 5).

**Figure 5:**
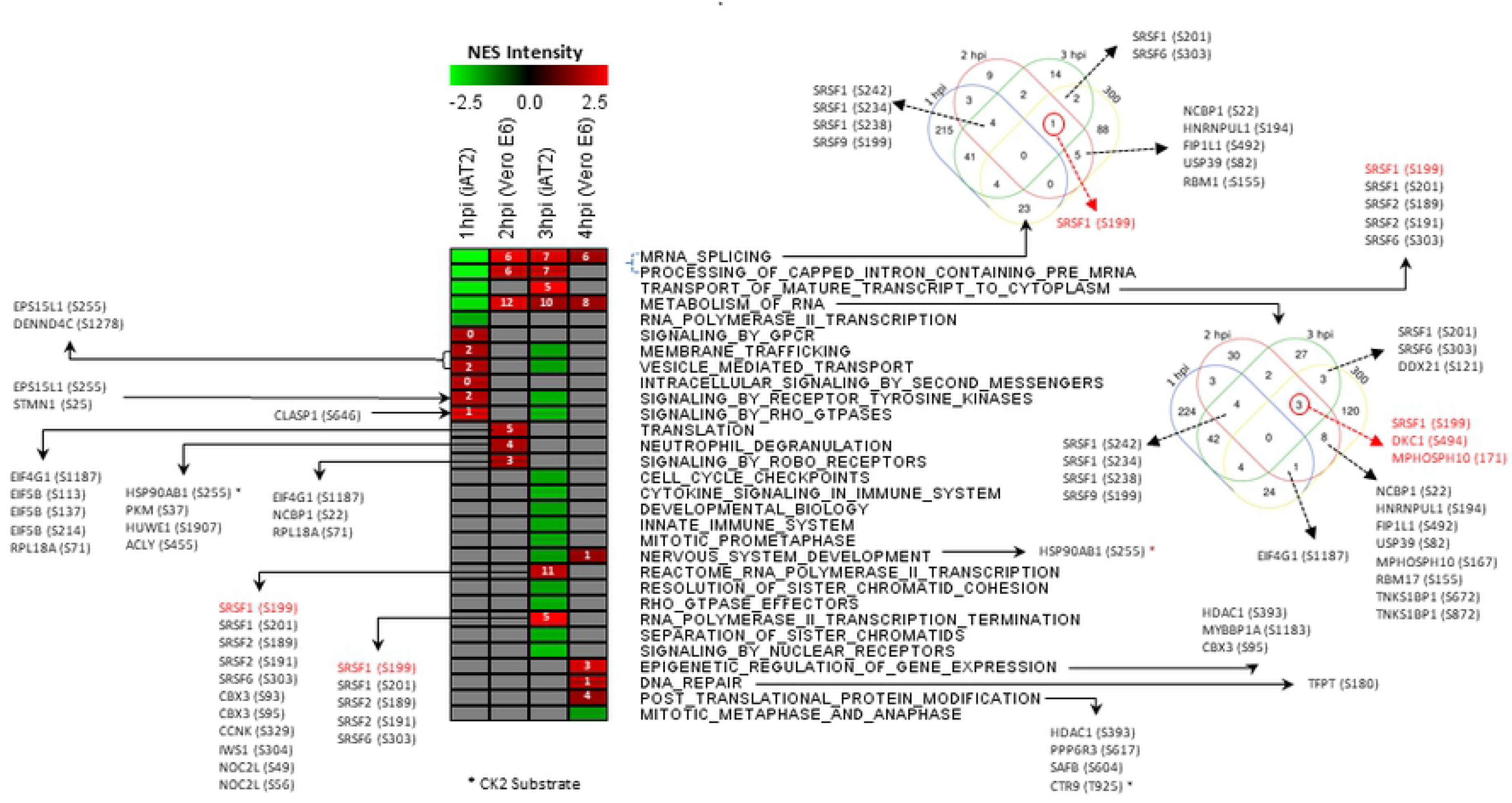
GSEA gene set enrichment analysis of Reactome pathways of proteins with altered phopsphorylation patterns in iAT2 and Vero cell lines. At the center of the figure is shown a heat map of the normalized enrichment scores resulted from GSEA analysis. The four columns corresponds to the 1h and 3h time points of iAT2 cell [15] and 2h and 4h time points of Vero cells [6] after infection by SARS-CoV-2. The numbers in small rectangles of the heat map indicate the number of proteins in the enriched set that contain phosphosites upregulated by SARS-CoV-2 infection and inhibited by CIGB-300 [9], these same sites are listed. The Venn diagram in the top right was built for the sets proteins of the “mRNA Splicing” Reactome pathways containing phosphosites that were differentially regulated at 1h, 2h and 3h time points or were inhibited by the action of CIGB-300 according to Perera et al. [9]. The second Venn diagram contains similar information for “Metabolism of RNA” Reactome pathway. All the sites upregulated at some time point by the infection and that were inhibited by CIGB-300 are listed.

SRSFs are RBP splicing factors that belong to the family of S/R rich proteins. Rogan et al. [50] proposed a molecular mechanism for viral-RNA pulmonary infections based on protein expression and RBP binding site pattern analysis. They compared the distribution of RBP binding motifs in several viral genomes including SARS_CoV-2, Influenza A, HIV-1 and Dengue. These authors identified strong RBPs binding sites in SARS-CoV-2 genome. After infection, as the number of SARS-CoV-2 genomes increase, the proportion of SRSFs bound to viral genome versus host transcriptome also increases. As the virus replicates in cytoplasm, newly synthetized SRSF1 molecules are bound by viral RNA and retained there, resulting in the formation of R-loops in the nucleus due to a reduction of RBP import. Rogan et al. [50] suggested that R-loop induced apoptosis could contribute to the spreading of viral particles to neighboring pneumocytes causing a deterioration of lung functions.

Phosphorylation plays an important role in SRSF proteins function. SRPK1 kinase was shown to phosphorylate multiple serine residues in SR rich domain of SRSF1 [51, 52], promoting its nuclear import where it plays an important role in RNA stability [53] and alternative splicing [54]. CK2 was found to be the major kinase that phosphorylate SRPK1 and this phosphorylation occurs mainly at Ser51 and Ser555, resulting in 6-fold activation of the enzyme [55]. After SARS-CoV-2 infection of AT2 cell, Ser51 is activated at 3h and 6h [15].

Fig 6 show the expression profile of CK2 and the levels of phosphorylation of SRPK1 S51 site, according to data from Hekman et al. [15]. A clear correlation is observed between the amount of CK2 kinase and the phosphorylation activation of this phospho-acceptor site, an additional argument supporting the role of CK2 on the activation of SRPK1 during SARS-CoV-2 infection.

**Figure 6:**
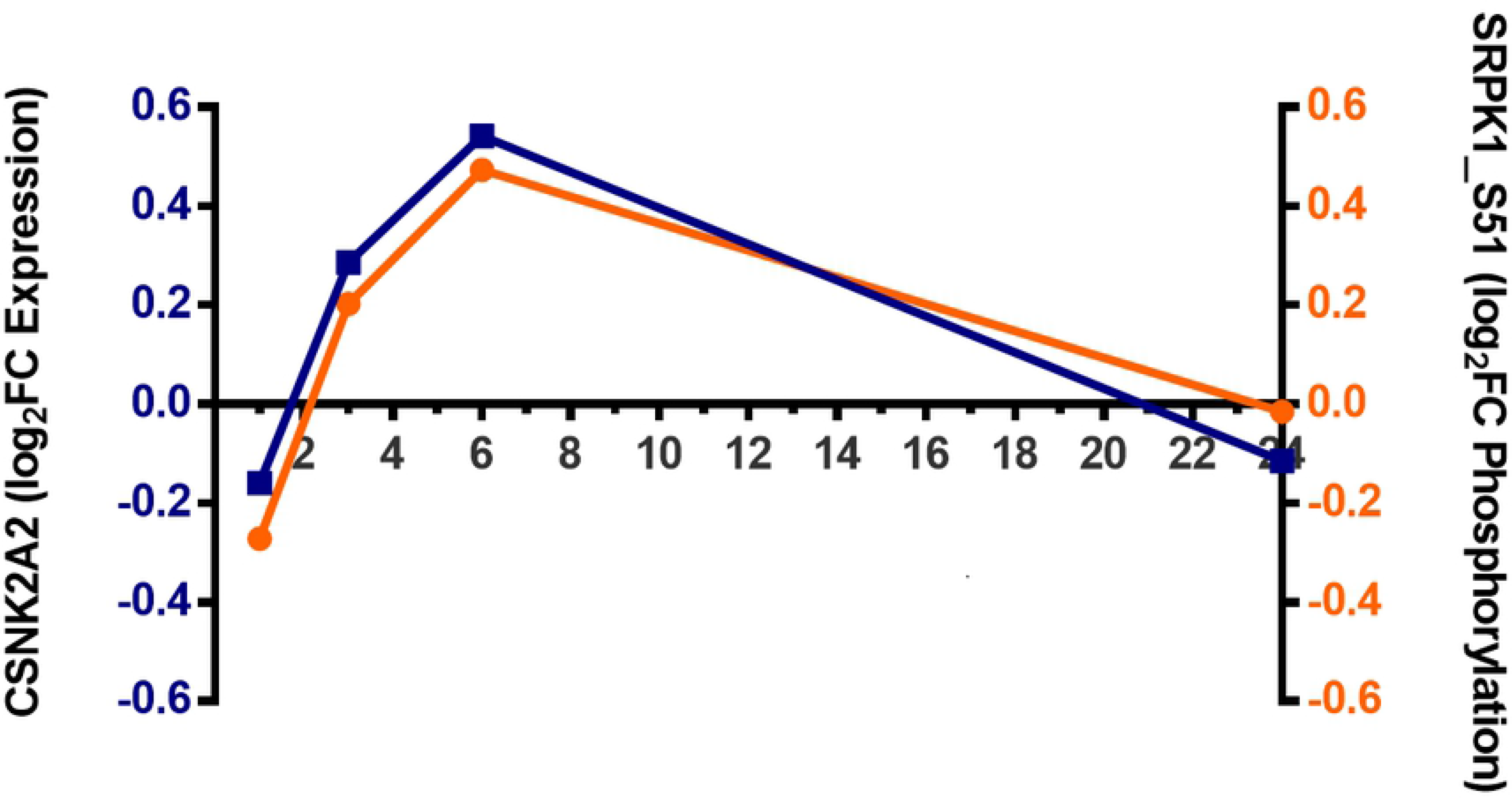
CK2 expression and SRKP1_S51 site phosphorylation profiles.

These results are consistent with previous reports predicting an extensive reshaping of splicing pathways by SARS-CoV-2 infection [16, 29]. SRSF1 is an important element of this splicing machinery that is clearly used by SARS-CoV-2 for its own replication and translation.

The increasing amount of SRSF1 bound to viral genome as the infection progress is a clear indication of its role in viral RNA processing. Phosphorylation is an important mechanism that control SRSF1 function. Taking all these together, we suggest that CIGB-300 intervene SRSF1 role in SARS-CoV-2 protein synthesis interfering its phosphorylation by SRPK1 kinase.

#### Infection-induced protein-protein interactions could be perturbed by CIGB-300

Next, we compared the host-viral PPIs reported by Gordon et al. [5] with phosphoproteomic data from Perera et al. [9] on the identification of CK2 substrates significantly inhibited by the CIGB-300. Fig 7 show virus-host interactions from Gordon et al. [5] in which host proteins contain phospho-acceptor sites that were inhibited by CIGB-300 treatment (highlighted in yellow). In this network several proteins are relate to RNA processing and transcription (LARP1, LARP7, LARP4B), supporting the results already mentioned. Binding of CIGB-300 to phospho-acceptor sites of host proteins inhibiting its phosphorylation may perturb the binding by viral proteins and consequently the viral life cycle.

**Figure 7:**
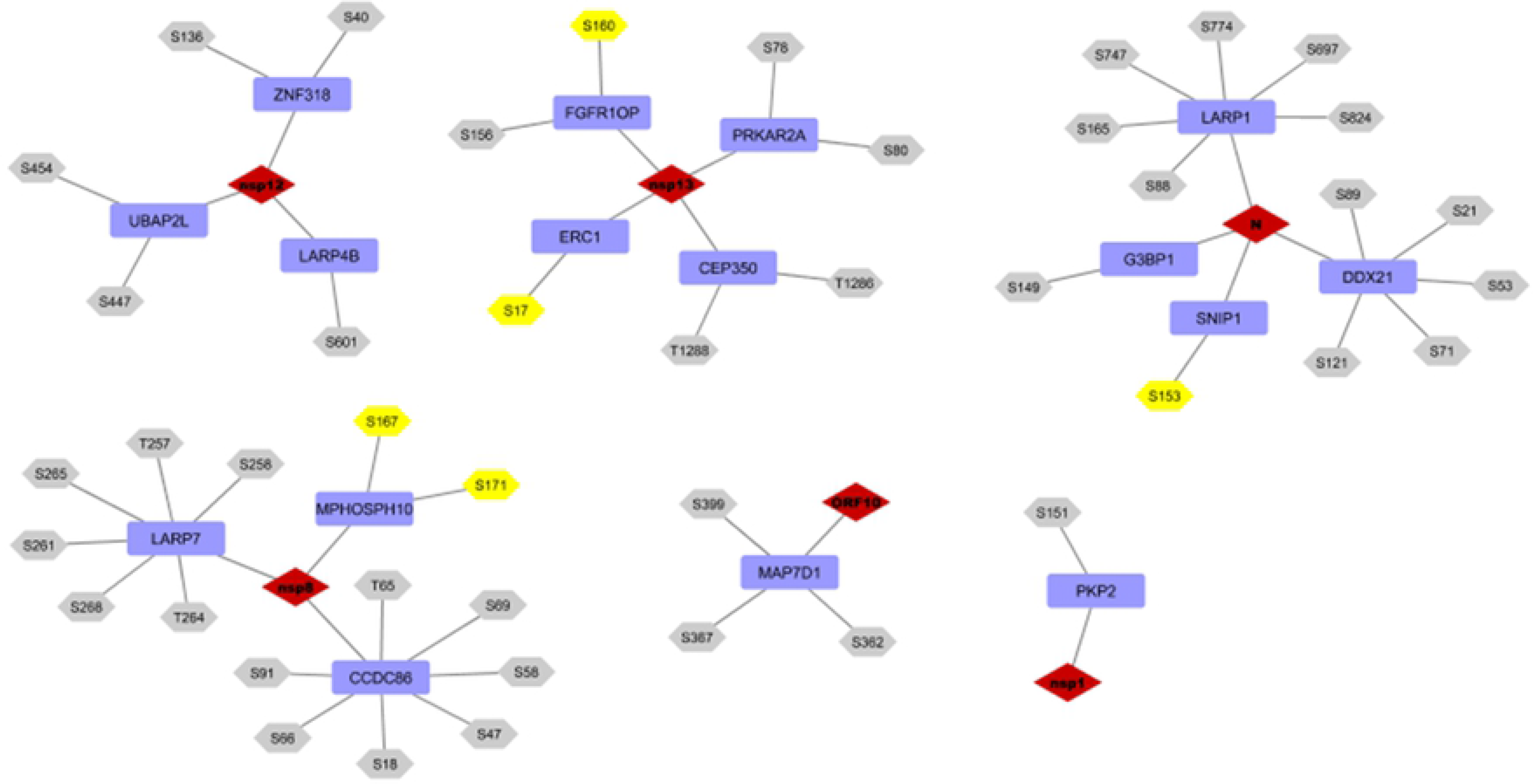
Viral protein interactions with host proteins with phosphorylation sites inhibited by CIGB-300. Rhombus in red represents viral proteins, rectangles in blue represent host proteins, hexagons represent phospho-acceptor sites. In yellow are shown those sites whose phosphorylation was increased by SARS-CoV-2 infection. Evaluating how CIGB-300 may interfere host-host protein interactions implicated in virus-induced mechanisms, we found that 68 proteins (SC2_300 set from now on) have activated phospho-acceptor sites in at least two of the four phosphoproteomic studies, which were inhibited by CIGB-300 (see S2 Table). The protein-protein interactions (PPI) network built with these proteins is shown in Fig 8. A majority of the nodes in the network are interconnected indicating potential functional relations among them of biological significance. Proteins are grouped by mRNA metabolism, Cell Cycle, and Selective Autophagy pathways, identified as significant by a Reactome enrichment analysis (see S3 Table). The five proteins with higher degree are also highlighted. Among them are HNRNPA1, HSPB1, SRRM2, and SRRM1, which are implicated in mRNA metabolism, corroborating the potential impact of CIGB-300 in viral replication and transcription. The fifth protein was B23/NPM1, identified as a major target of CIGB-300 in cancer cells, but also as a relevant target for antiviral therapies [10–12].

In this network HSPB1 heat shock protein (alias HSP27) is one of the highest degree nodes. HSPB1 was found to be overexpressed in idiopathic pulmonary fibrosis (IPF) patients. It activates pro-fibrotic mechanisms and consequently has been suggested as a target to treat IPF [56, 57]. In tumor cells, Ivermectin inhibits the phosphorylation of Ser78 and Ser82 of HSP27, while Ser15 is only slightly inhibited [58]. Also Ivermectin have shown to be on inhibitor of SARS-CoV-2 with a significant reduction of viral RNA levels [59] and increase the clinical recovery of mild and severe Covid-19 patients [60, 61]. SARS-CoV-2 activates HSPB1 Ser15 and Ser82 during infection while CIGB-300 inhibits both phospho-acceptor sites [9]. This is an additional argument in favor of using CIGB-300 in Covid-19 patients aiming to reduce pulmonary lesions as it was evidenced in a phase I/II clinical trial [13].

### Human phenotypes involving kinase activity induced by SARS-CoV-2, potentially targeted by CIGB-300

We build a network with the top 20 human phenotypes most enriched in the set SC2_300, using GeneCodis tool (see Fig 9 and S4 Table). The network can be divided in two main subnetworks, one related to muscular disorder phenotypes that include Paralysis, Distal muscle weakness, Rimmed Vacuoles, Mildly elevated creatine kinase and Fatigue. The second subnetwork groups phenotypes related to respiratory(Exertional dyspnea, difuse alveolar hemorraghe), bleeding (Metrorrhagia, oral cavity bleeding) and coagulation disorders(Disseminated intravascular coagulation).

**Figure 8:**
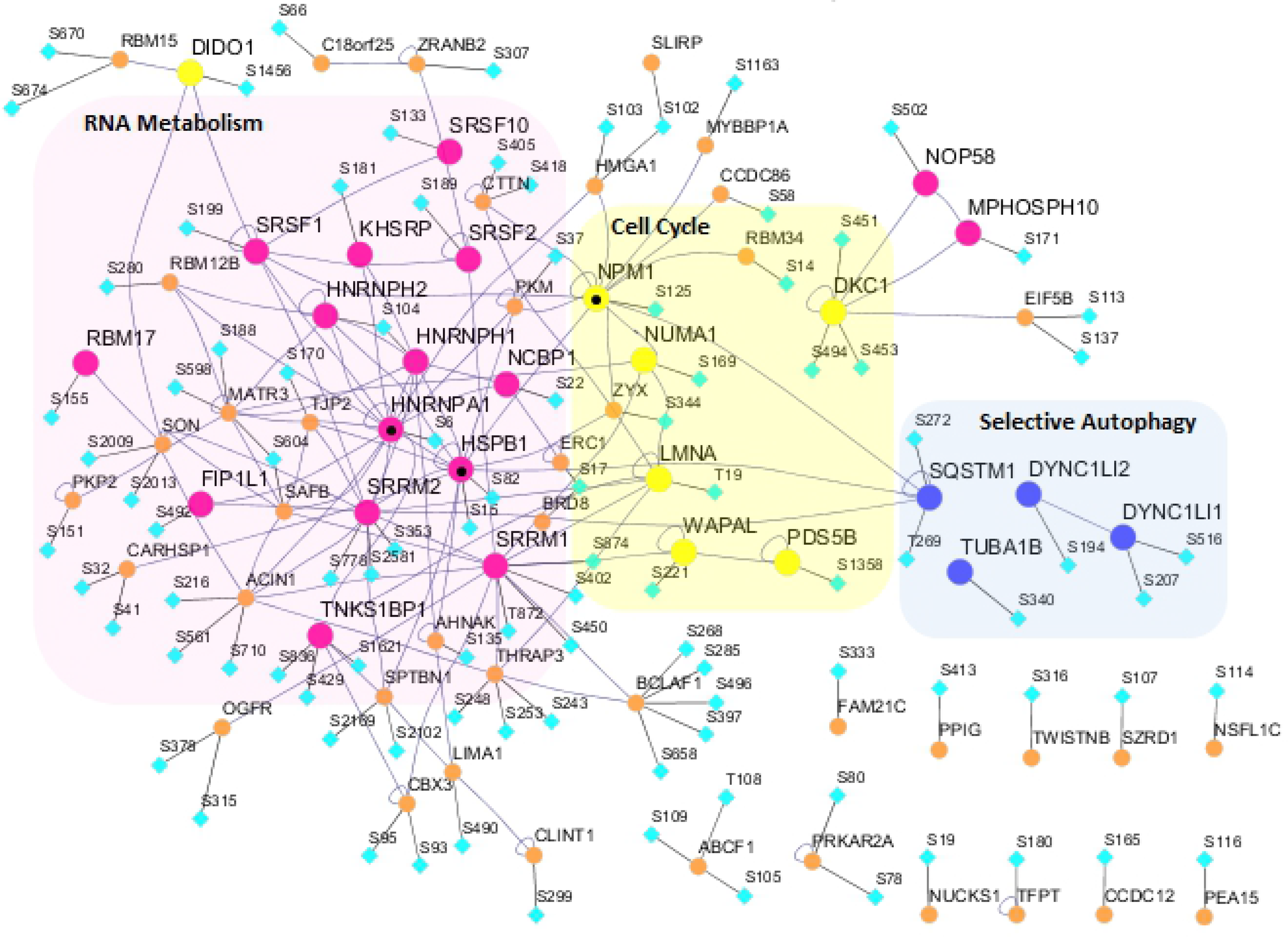
Interaction network of proteins found to be activated in at least two of the four phosphoproteimics studies analyzed and with phosphoaceptor sites found to be inhibited by the action of CIGB-300. For each protein the phospho-acceptor sites inhibited by CIGB-300 are shown. Proteins involved in more significant pathway are grouped and colored: mRNA metabolism(•), Cell Cycle(•) and ‘Selective Autophagy’ (•). The five nodes with a higher degree (HNRNPA1, HSPB1, SRRM2, NPM1 and SRRM1) are labeled with (•).

**Figure 9:**
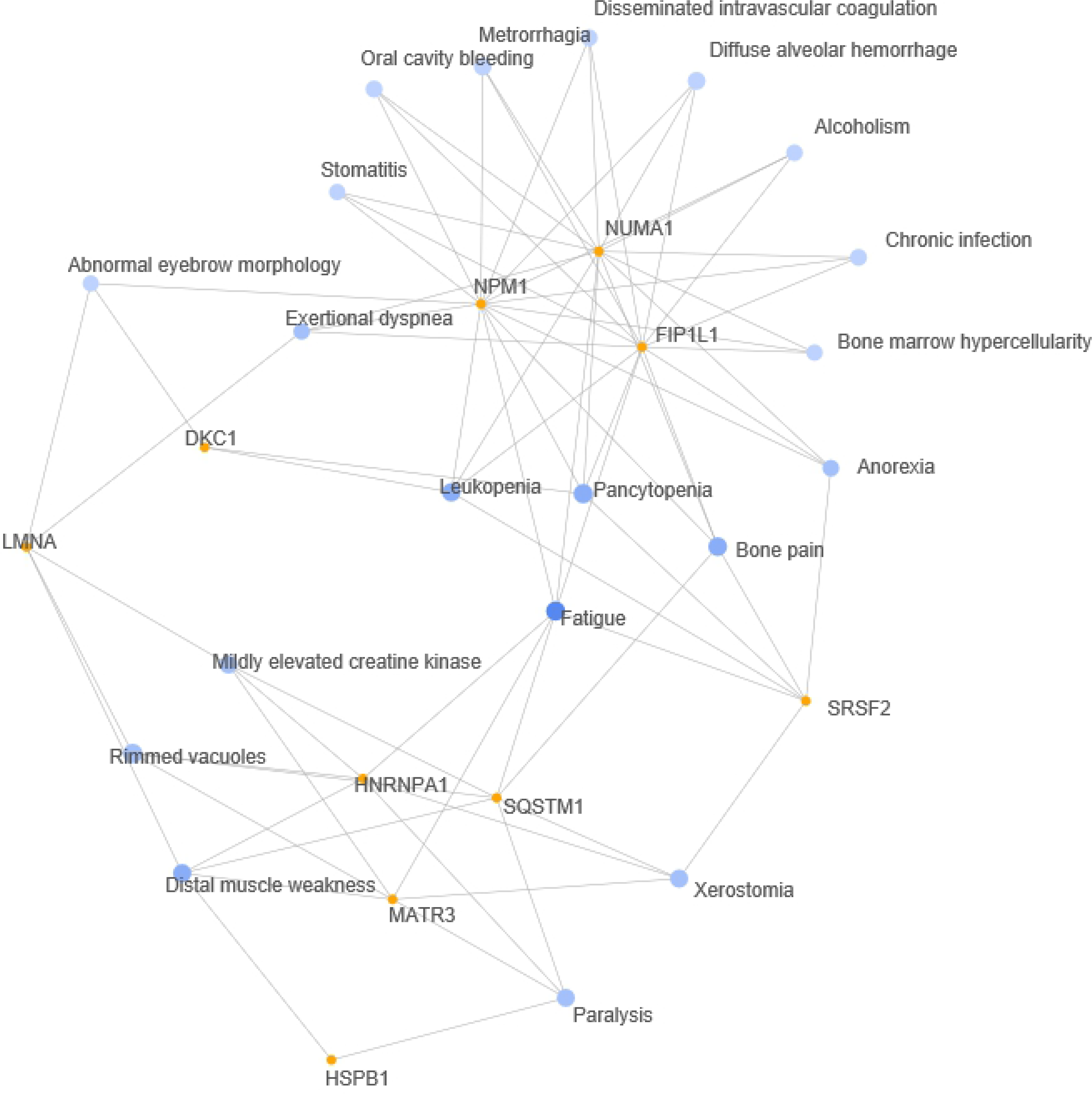
Network representation of Human Phenotypes (HPO) enriched in a set of 68 proteins that contain phospho-acceptor-sites that were activated by SARS-CoV-2 infection in at least two of the four phosphoproteomics studies and inhibited by the action of CIGB-300. Genecodis functional annotation tool was used for the enrichment analysis and to generate the network of the top 20 enriched phenotypes. Nodes in blue represents enriched phenotypes and nodes in orange represent proteins associated to these phenotypes.

The phenotypes in the first subnetwork are all associated to HNRNPA1, MATR3 and SQSTM1 genes, and have been also reported as Covid-19 symptoms [62–66]. For example, elevated creatine kinase levels is associated to a poor outcome prediction [62] and persistent fatigue is a common symptom in Covid-19 patients [65]. As we mentioned MATR3 and SQSTM1 possess phosphosites that were activated in all four phosphoproteomic studies we analyzed, and HNRNPA1 was the node with the highest degree in the PPI network built. Rimmed Vacuoles, the most significant of the enriched phenotypes, are found in areas of destruction of muscle fibers. Fatigue phenotype is located somewhere in the interface between the two subnetworks and is connected to the three genes mentioned, and also to the three genes that are in the core of the second subnetwork: B23/NPM1, FIP1L1 and NUMA1. The phenotypes in the second subnetwork have been all identified in Covid-19 patients [67, 68].

### Targeting B23 chaperone activity by CIGB-300

Again B23 surfaced as a relevant player of a viral infection, now in the context of SARS-CoV-2. First, we saw it as the host protein with a higher correlation of expression to viral proteins, in particular to Orf6. Second, we identified B23 as a highly connected node in a network of proteins consistently upregulated by SARS-CoV-2 infection and inhibited by CIGB-300, which is related to Cell Cycle pathway (Fig 8). Third, it was part of a phenotype network related to respiratory, bleeding and coagulation disorders, symptoms widely reported in Covid-19 disease. Previously, Kondo et al. [69] showed that B23 inhibited the DNA-binding and transcriptional activity of interferon regulatory factor 1(IRF1), while Abe et al. [70] found that B23 regulates the expression of IFN-γ-inducible genes and binds to transcription factors STAT1 and IRF1. Taken together, both Orf6 and B23 might play a role in the inhibitory effect of IFN signaling.

On the other hand, it is known that post-translational modifications like phosphorylation are involved in the regulation of molecular chaperone activities [71]. In particular CK2 phosphorylation was found to play an important role in B23 chaperon activity [72].

CIGB-300 might interfere B23 chaperon activity by inhibiting phosphorylation and perturbing its interactions with host and viral proteins. For instance, Orf6 localize in ER/Golgi membrane and NPC associated to KPNA2 and NUP98, respectively. May B23 chaperone activity play a role in SARS-CoV-2 infected cells by carrying Orf6 to the ER/Golgi membrane and the NPC?

Of note, the CIGB-300 peptide interacted with the B23 protein in MDKB cells infected with a Bovine Coronavirus strain (BCoV) [14]. In these cells, also several host proteins participating in protein folding, populated the interactomic profile of the CIGB-300 peptide. However, to definitively address any particular role of B23/NPM1 in the context of an ongoing coronavirus infection, gain- and/or lost-of-function genetic experiments need to be done.

## Conclusion

There is, so far, no definite effective therapeutic treatment against SARS-CoV2, despite progress with the development and extensive use of vaccines. The emergence of increasingly transmissible and aggressive mutated variants of the virus justifies the efforts to find new therapeutics to reduce the ratio of patients that evolve to severe stages and death.

Targeting those mechanisms of host cell that are commonly hijacked by viruses, to reproduce and spread, is a valid strategy to confront present and future challenges of viral epidemics. CK2 kinase activity is activated by multiple viruses and proved to be essential for many virus-induced mechanisms that allow viruses to replicate and damage host cell functions. Phosphorylation is widely altered in human cell immediately after virus entry, contributing to the hijacking of multiple cellular processes.

In this paper putative molecular mechanism are put forward to support the observed antiviral effects of CIGB-300 in coronavirus infections and the preliminary clinical results on preventing virus induced lung injury.

Our *in-silico* analysis provides evidences of the role of CIGB-300 in the reversion, based in CK2 kinase activity inhibition, of virus-induced perturbations of cellular functions such as:

- N protein localization and RNA binding capabilities which are implicated in essential steps of viral life cycle as viral capside assembly.
- Interference of IFN signaling, which is responsible for a weak immune response.
- Early stages of virus kidnapping of RNA processing, which plays an essential role in virus genome replication and translation.
- Aggresome accumulation and its role in Inflammation and fibrosis commonly observed in severe Covid-19 patients.
- Phenotypes related to muscle, bleeding, coagulation and respiratory disorders.

Table 1 summarizes different working hypothesis which need to be verified in suitable pre-clinical models. Notably, while our findings suggest a clear impact of CK2 inhibitors in viral replication and hijacking strategies, differences could be also inferred based on their particular inhibitory mechanism. For instance, the use of CIGB-300 to impair CK2-mediated signaling in cancer do not mirror CX-4945 effects in pre-clinical and clinical settings, thus the same could be expected in viral infections like those caused by coronaviruses. The fact that CIGB-300 targets both the CK2 enzyme and a subset of its substrates, may imply particular inhibitory effects of protein-protein interactions, as well as in the crosstalks with other nearby Post-translational modifications sites [9]; therefore, resulting into different molecular, cellular and organismal outcomes.

**Table 1:** Summary of main findings.

| Analysis Type | Subject | Identity | Working Hypothesis | Experimental Clues | Reference |
| --- | --- | --- | --- | --- | --- |
| Individual CK2 sites | Viral proteins | N | CK2 phosphosite inhibition/blockage by CIGB-300 impairs viral replication and IFN signaling | Interaction, co-localization, N mRNA and protein expression inhibition by CIGB-300 in a subrogate model | [14] |
| Individual CK2 sites | Viral proteins | ORF6 | CK2 phosphosite inhibition/blockage by CIGB-300 impairs IFN signaling | Additive/Synergistic profile of CIGB-300 plus IFN alpha | Unpublished |
| Co-Expression and network propagation | Host and Viral Proteins | NUP98, ORF6, B23 | CK2 phosphosite inhibition/blockage by CIGB-300 impairs IFN signaling | Additive/Synergistic profile of CIGB-300 plus IFN alpha (unpublished) | Unpublished |
| Phosphoproteome overlap | Host Proteins | MATR3, SQSTM1, DIDO1 | CIGB-300 Multitarget effect impairing viral transcription/splicing, inflammation, immunoresponse and apoptosis | Pulmonary lesions resolution in CT Phase I | [13] |
| Kinase activity profiles | Host proteins | SRPK1 (SRSF1,2,6,10) | CIGB-300 impairs Viral RNA splicing | None, to be evaluated in preclinical settings | NA |
| PPIs vs H125 Phosphoproteome | Host and Viral Proteins | Several proteins (see text and Fig. 7)<br>[LARP1,LARP7,LARP4B] | CIGB-300 impairs Viral RNA processing and transcription | None, to be evaluated in preclinical settings | NA |
| SARS-CoV-2 vs H125 Phosphoproteome overlap, Network and Enrichment analysis | Host proteins | Several proteins (see text and Fig. 8)<br>[HNRNPA1, HSPB1, SRRM2, SRRM1, B23] | CIGB-300 Multitarget effect impairing on mRNA metabolism cell cycle and Autophagy pathways | Pulmonary lesions resolution in CT Phase I | [13] |
| Human Phenotypes | Host proteins | HNPRNPA1, MATR3, SQSTM1, B23, FIP1L1, NUMA1 | CIGB-300 might relief COVID19 clinical symptoms | To be evaluated in CT Phase II | NA |

Altogether, our computational biology approach supports the potential impact of CIGB-300 in the treatment of Covid-19 patients, and encourages its clinical evaluation from early stages of the viral infection.

## Material and Methods

### Public Data

Protein sequence information was downloaded from UniprotKB database [73] https://www.uniprot.org/.

Data sources from phosphoproteomics studies of host and viral proteins of SARS-CoV-2 infected human cell lines were downloaded from site listed in table 2.

**Table 2:** SARS-CoV-2 infection phosphoproteomics studies data sources.

| Reference | Timepoints | Cell lines | Downloaded from: |
| --- | --- | --- | --- |
| Bouhaddou et al (2020) | 2h,4h,8h,12h and 24h | Vero E6 | http://dx.doi.org/10.17632/dpkbh2g9hy.1 |
| Hekman et al (2020) | 1h,3h,6h and 24h | iAT2 | https://www.ebi.ac.uk/pride/archive/projects/PXD020183 |
| Stukalov et al (2021) | 6h and 24h | A549 | https://www.nature.com/articles/s41586-021-03493-4#MOESM5 |
| Klann et. al (2020) | 24h | CaCo-2 | [https://www.cell.com/molecular-cell/fulltext/S1097-2765(20)30549-9#supplementaryMaterial](https://www.cell.com/molecular-cell/fulltext/S1097-2765%2820%2930549-9#supplementaryMaterial) |

The phosphorylation data of CIGB-300 treatment from Perera et al. [9] were downloaded from https://link.springer.com/article/10.1007%2Fs11010-020-03747-1#additional-information/.

Protein expression data from Bojkova et al. [29] were downloaded from http://corona.papers.biochem2.com/.

A total of 332 humand-SARS-CoV-2 protein-protein interactions from Gordon et al. [5] were downloaded from Biogrid database [74] at https://thebiogrid.org/225737/publication/comparative-host-coronavirus-protein-interaction-networks-reveal-pan-viral-disease-mechanisms.html#!.

Information of CK2 Substrates was downloaded from PhosphoSitePlus at (https://www.phosphosite.org/) [75].

### Data Analysis

Sequence alignments were performed with the desktop version of CLUSTALX multiple sequence alignment program [76].

Analysis of proteomics data from Bojkova et al. [29] was carried out with MEV [77]. SAM method [78] for two-class unpaired comparison was used to identify most differentially expressed proteins as implemented in MEV. Parameter delta was set so that the estimated median number of false significant proteins would be cero.

For Bouhaddou et al. [6], Stukalov et al. [17], Klann et al. [16] and Hekman et al. [15] data on phosphorylation induced by SARS-CoV-2 infection, the criteria for choosing differentially phosphorylated sites was log_2_ fold change>=0.25 and adjusted-p-value<=0.05, the same values proposed by Hekman et al. [15].

The phosphorylation changes induced by CIGB-300 treatment were extracted from Perera et al. [9]. All inhibitions reported by the authors at 10 and/or 30 minutes were considered.

For Gene Set Enrichment Analysis we used GSEA version 4.1.0 for windows [79, 80]. Ordered gene lists for each time point after infection were provided as input. The pre-ranked gene list option was used for databases containing REACTOME and GO biological process gene sets. MSigdb v7.2 gmt files were downloaded from: http://www.gsea-msigdb.org/gsea/downloads.jsp.

Cytoscape tool [81, 82] was used to build and merge networks in Figs 7 and 8.

BisoGenet CytoScape plugin [83], available from CytoScape Application Manager, was used to generate Protein-protein interaction (PPI) networks.

Venn Diagrams were generated using web application in http://bioinformatics.psb.ugent.be/.

Functional analysis of enriched pathways and reactions was performed using Reactome Pathway Knowledgebase [84] at: https://reactome.org/. The criteria used for selection was FDR (False Discovery Rate) <=0.05.

GeneCodis 4.0, at https://genecodis.genyo.es/, was used for diseases enrichment analysis [85]. Data sources for Human Phenotypes HPO and OMIM were both consulted individually and integrated. The results were obtained in form of networks clusters.

BiNGO plugin [86], available from Cytoscape Application Manager, was used to determine and visualize Gene Ontology (GO) categories statistically overrepresented.

Additional statistical analysis and graphs were generated and plotted using GraphPad Prism version 5.00 software (GraphPad Software, San Diego, CA, USA).

## Acknowledgments

We thank Ricardo Javier Bringas for the design and drawing of figure 2.

## Author Contributions

**Conceptualization:** Jamilet Miranda, Ricardo Bringas, Yasser Perera.

**Data curation:** Jamilet Miranda, Ricardo Bringas.

**Formal analysis:** Jamilet Miranda, Ricardo Bringas.

**Investigation:** Jamilet Miranda, Ricardo Bringas, Yasser Perera.

**Methodology:** Jamilet Miranda, Ricardo Bringas, Jorge Fernandez de Cossio, Yasser Perera.

**Supervision:** Ricardo Bringas, Yasser Perera.

**Writing – original draft:** Jamilet Miranda, Ricardo Bringas.

**Writing – review & editing:** Jamilet Miranda, Ricardo Bringas, Jorge Fernandez de Cossio, Yasser Perera.

## Supporting information

**S1 Fig. Bidimensional clustering of most differentially expressed proteins resulting from the analysis of proteomic data of CaCo-2 cells infected by SARS-CoV-2 from Bojkova et al.(2020).** Horizontal axis represents samples at different time points and vertical axis represent differentially expressed proteins. Rows corresponding to viral Orf6 and host B23 proteins are labeled.

**S1 Table. N phospho-acceptor sites reported in four phosphoproteomics studies*.**

**S2 Table. List of 102 phosphosites activated in at least two of the four phosphoproteomic studies.** Level 1 (in 4 experiments), level 2 (in 3 experiments) and level 3 (in 2 experiments).

**S3 Table. REACTOME pathways enrichment results.**

**S4 Table. Human phenotypes enrichment results.**

